# Linked selection and the evolution of altruism in family-structured populations

**DOI:** 10.1101/2022.05.15.492036

**Authors:** Lia Thomson, Daniel Priego Espinosa, Yaniv Brandvain, Jeremy Van Cleve

## Abstract

Much research on the evolution of altruism via kin selection, group selection, and reciprocity focuses on the role of a single locus or quantitative trait. Very few studies have explored how linked selection, or selection at loci neighboring an altruism locus, impacts the evolution of altruism. While linked selection can decrease the efficacy of selection at neighboring loci, it might have other effects including promoting selection for altruism by increasing relatedness in regions of low recombination. Here, we used population genetic simulations to study how negative selection at linked loci, or background selection, affects the evolution of altruism. When altruism occurs between full siblings, we found that background selection interfered with selection on the altruistic allele, increasing its fixation probability when the altruistic allele was disfavored and reducing its fixation when the allele was favored. In other words, background selection has the same effect on altruistic genes in family-structured populations as it does on other, nonsocial, genes. This contrasts with prior research showing that linked selective sweeps can favor the evolution of cooperation, and we discuss possibilities for resolving these contrasting results.

## Introduction

Altruism is the phenomenon where an organism improves another’s fitness at the expense of its own fitness (Hamilton, 1964; Lehmann & Keller, 2006; Rousset, 2004; Taylor, 1992). This type of behavior is found in many species, from microbes like *Dictyostelium discoideum* (Noh et al., 2018; Strassmann et al., 2000) to aphids (Benton & Foster, 1992; Uematsu et al., 2010), ants (Bourke & Franks, 1995; Hamilton, 1972), and primates including humans (Burkart et al., 2007; Silk & Boyd, 2010; Warneken et al., 2007). The existence of altruism despite its costs to survival or reproduction has spawned a tremendous amount of evolutionary theory. Multiple explanations have been proposed, such as kin selection and reciprocal altruism or reciprocity (Axelrod & Hamilton, 1981; Hamilton, 1964; Maynard Smith, 1974; Trivers, 1971). The foundational research on the evolution of altruism occurred squarely in the “pre-genomic” era — before the evolutionary consequences of selection at loci genetically linked to the locus of interest (e.g., Charlesworth et al., 1993) were seriously considered. Since then, the immense theoretical (Akçay, 2018; Akçay & Van Cleve, 2012; Allen et al., 2017; e.g., Hamilton, 1970; Lehmann & Keller, 2006; McAvoy et al., 2020; Ohtsuki et al., 2006; Queller, 1985; Rankin, 2011; Rousset & Billiard, 2000; Tarnita et al., 2009; Taylor & Frank, 1996; Úbeda & Gardner, 2010; Van Cleve, 2015, 2017, 2020) and empirical (Boomsma, 2009; Bourke, 2014; Griesser et al., 2017; Krakauer, 2005; Lukas & Clutton-Brock, 2012; Nadell et al., 2016; Ostrowski, 2019; Strassmann et al., 2011) progress on the evolution of altruism has largely ignored the potential influence of selection on linked genes whose expression is not related to cooperation or social behavior.

In general, selection on non-neutral mutations generates a stochastic evolutionary force that decreases genetic diversity at linked sites (i.e., linked loci). This can be seen in background selection (Charlesworth et al., 1993; Hudson & Kaplan, 1995), which is caused by linked genes with small deleterious effects, or in a selective sweep (Kaplan et al., 1989; Maynard Smith & Haigh, 1974), which is caused by a linked gene with a strong beneficial effect. Linked selection is most influential in genomic regions with low recombination rates. Another factor affecting genetic diversity is spatial structure and limited migration between local populations or demes (Crow & Kimura, 1970): when migration among demes is weak, individuals tend to reproduce in the same deme as their parents, and they are more likely to have identical alleles than individuals in different demes. This results in a decrease in local genetic diversity within demes and an increase in genetic relatedness as measured by the *F*_ST_ statistic (Wright, 1931, 1951), which compares genetic identity within demes to genetic identity between demes. In spatially structured populations, the effect of linked selection is to increase genetic homogeneity among linked loci within demes as measured by an increased *F*_ST_ (Charlesworth et al., 1997; Hu & He, 2005; Slatkin & Wiehe, 1998, 1998).

Genetic relatedness is also one of the key ingredients in the evolution of altruism. Hamilton (1964) showed that the total effect of natural selection on an altruistic allele (Lehmann & Rousset, 2014; assuming additive genetic effects and weak selection; Rousset & Billiard, 2000) is given by the inclusive fitness effect: –*c* + *r b* where *c* is the fitness cost to the helping individual, *b* is the fitness benefit to the individual helped, and *r* is the genetic relatedness between the two (Hamilton, 1964), which can be measured by *F*_ST_ (Hamilton, 1964, 1970; Rousset, 2004). Hamilton’s rule states that an altruistic allele is favored by selection when *c/b* < *r* (i.e. the inclusive fitness effect is greater than zero). Because linked selection within a genomic region may increase genetic relatedness in that region, Hamilton’s rule suggests that it might also increase the range of cost/benefit ratios where an altruistic allele in that genomic region may evolve by natural selection. In fact, both theoretical and empirical research in asexual microbial systems suggest that selective sweeps caused by linked beneficial mutations unrelated to the social interaction can promote the evolution of altruism (Hammarlund et al., 2016) and protect communities from invasion by “selfish” cheaters (Morgan et al., 2012). However, these complex models and experiments in microbial systems combined various ecological and evolutionary processes (e.g. extinction and recolonization, fluctuating population sizes, etc.) and therefore it is unclear if a change in relatedness due to linked selection or some other feature of the biology of these systems is responsible for the increased benefit of cooperation in these cases.

To test the hypothesis – implied by results from microbial systems – that linked selection itself enhances the evolution of altruism, we conducted simulations of family-structured populations of sexual, recombining diploids. Family-structured populations, which were the original context for Hamilton’s theory of kin selection (Hamilton, 1964), limit social interactions to within families and allow us to isolate the effect of linked selection via background selection on altruism since they generate simple baseline relatedness values without invoking population structure where background selection is known to have a complex relationship with relatedness via *F*_ST_ (Matthey-Doret & Whitlock, 2019). We show that rather than increasing the range of cost/benefit ratios (*c/b*) under which an altruism is favored, background selection simply weakens the efficacy of natural selection, increasing (or decreasing) fixation probabilities when the altruistic allele was disfavored (or favored. Thus, in contrast to expectations from selective sweeps in asexual microbial systems, background selection affects social traits within families as it does all other traits – by increasing the power of drift and decreasing the efficacy of selection.

## Methods

We wrote and executed all simulations in SLiM 3 (Haller & Messer, 2019). We used the “non-Wright-Fisher” (nonWF) mode in SLiM (see “Data Availability Statement” for SLiM code) since it allows for fitness to be affected by social interactions among individuals. In nonWF simulations in SLiM, most demographic events in the lifecycle are specified explicitly including density dependent regulation of population size, which occurs in our simulation through a carrying capacity parameter that we set equal to 1000 individuals. This leads to an equilibrium population size very close to 1000 with small deviations below that value due to the deleterious mutation load. Juveniles mature into adults after density dependent regulation and adults are killed off after reproducing, which leads to non-overlapping generations. Each individual is diploid with a 100kb-long genome. A single locus that can have alleles that cause altruistic behavior is located at base pair 1000. Recombination occurred along the genome at a rate of *ρ* = 10^−8^ per base pair. To represent background selection, slightly detrimental mutations occur at a rate of *u* = 5×10^−8^ per base pair in the whole 100kb region. Across simulations, we modulated the strength of background selection by varying the fitness effect of deleterious mutations from *s* = –0.006 to *s* = –0.014 in 0.002 intervals (dominance coefficient fixed at *h* = 0.5). For simulations without background selection, we used both neutral and strongly deleterious mutations by setting *s* = 0 and *s* = –1, respectively; neutral mutations do not affect fitness and cannot cause linked selection and strongly deleterious mutations are often removed by selection before they can have an appreciable effect on the evolutionary dynamics of the chromosomes on which they occur (e.g., they have a very small effect on the effective population size *N*_e_ and cause very little background selection; see equation 2 below).

For time efficiency, ten seed populations were generated by running the simulation for 5000 generations without the altruistic allele, which was sufficient to reach mutation-selection balance for deleterious variants (Supplementary Figure S1). All simulations start by loading these seed populations. Then, one copy of the altruistic allele is inserted into a random chromosome at base-pair 1000. This occurs after offspring are created for the generation, but before individuals die, such that the population size is larger than the carrying capacity, including roughly 1000 parents and 2000 offspring. Accordingly, the initial frequency and the fixation probability of a neutral allele inserted by this method is 1/6000.

During the reproduction phase of each generation, every member of the population is paired with another, if possible, and they produce a litter of four full siblings. Within the litter, one pair of siblings is randomly selected to engage in a social interaction that will affect their probability of survival to adulthood; the survival probability of the remaining two siblings who do not engage socially is unchanged. We assume the altruistic allele is completely dominant, so each sibling in the social interaction is altruistic to its partner if it has at least one copy of the altruistic allele. Performing an altruistic act decreases survival probability by a cost, *c*, and having an altruistic partner increases survival probability by a benefit, *b*. Thus, interacting pairs of altruistic siblings increase their survival probability by *b*–*c*. Unless otherwise noted, the cost of helping was kept at *c* = 0.1. We varied the benefit *b* across a range of cost/benefit ratios where altruism was favored and disfavored according to Hamilton’s rule. One million replicate simulations were run for each parameter combination.

We compared simulation results to fixation probabilities from population genetic theory. For the two cases without background selection (neutral linked mutations with *s* = 0 and strong deleterious linked mutations with *s* = –1), the fixation probability of an allele with dominance coefficient *h* and selection coefficient *𝒥* in a diploid population of effective size *N*_e_ is (Ewens, 2004, eqn. 3.30)

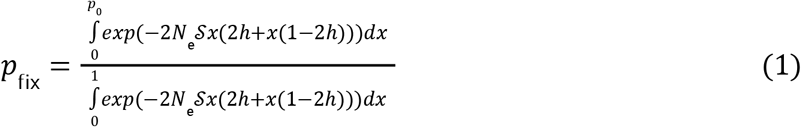

where *p*_0_ is the initial frequency of the allele. Since the family-structured model in our SLiM simulations is not an exact Wright-Fisher model, we needed to calculate the appropriate effective population size *N*_e_. In particular, the family structure and sampling created a different variance in reproductive success, σ^2^, than is typical for a Wright-Fisher model. We measured the variance in reproductive success with a SLiM script that tracked for each chromosome the number of copies of a locus at base pair 1000 that survive into the next generation. Without background selection, we calculated the variance in reproductive success in our simulation as σ^2^ ≈ 0.75. The effective size that accounts for the variance in reproductive success is *N*_e_ ≈ *N*/σ^2^ (Ewens, 2004, eqn. 3.107), which yields *N*_e_ ≈ 1333. The selection coefficient *𝒥* for the altruistic allele can be derived from the expected change in allele frequency and is approximately the inclusive fitness effect – *c* + *r b* (Rousset & Billiard, 2000; Roze & Rousset, 2004). In our simulations, since only one of the two pairs of siblings have the opportunity for altruism, our selection coefficient is one half the inclusive fitness effect: *𝒥* = (– *c* + *r b*)/2 = (– *c* + 0.5 *b*)/2. Using this expression for *𝒥, h* = 1, *p*_0_ = 1/6000, and *N*_e_ = 1333, we integrated equation (1) in Mathematica (Wolfram Research, Inc., 2021) to generate the predicted fixation probabilities in Figures 1 and S2. The vertical black lines represent the 95% binomial confidence intervals around the predicted fixation probabilities.

**Figure 1.**
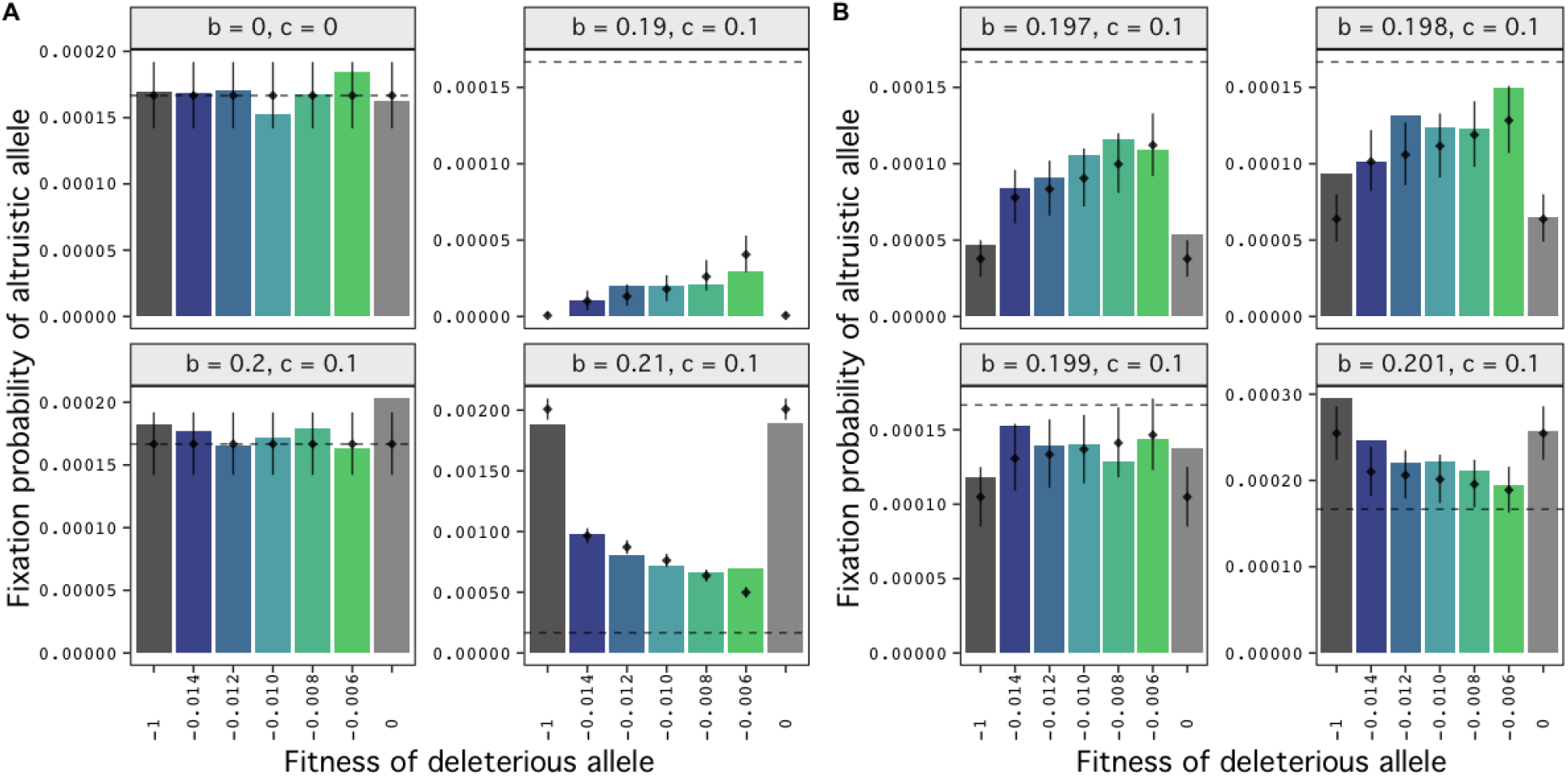
The effect of background selection on the fixation probability of the altruistic allele depends on whether the allele is selected for by Hamilton’s rule. Simulations of altruism between full siblings (*r* = 1/2) are plotted by the cost (*c*) and benefit (*b*) to fitness of an altruistic interaction. Shown in A) are two scenarios when the altruistic allele is expected to be neutral by Hamilton’s rule (leftmost column), a scenario when it is expected to be detrimental (upper right), and a scenario when it is expected to be beneficial (lower right). In B) benefit is varied by smaller increments around the expected neutral value of 0.2. The horizontal dashed line is the neutral fixation probability, and vertical lines indicate the 95% binomial confidence interval for the predicted fixation probabilities (black dots)incorporating Hudson and Kaplan’s (1995) approximation of the effect of background selection on *N*_e_ from equation (2) (see Methods for details).

For the cases with background selection (linked mutations with –0.014 ≤ *s* ≤ –0.006 in our simulation), we predicted the fixation probabilities of the altruistic allele assuming that the only effect of background selection is a change in the effective population size *N*_e_ in equation (1), which we call *N*_e,bg_. The expression for *N*_e,bg_ in this case is derived from equation (7)in Hudson and Kaplan (1995) by setting their *L*_1_ = 0 and *L*_2_ = *l* (since the altruistic allele in the simulation is on one end of the chromosome) and obtaining

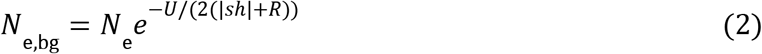

where *U* is the total diploid rate of deleterious mutation in the genomic region, *R* is the length of the genomic region in Morgans, and *N*_e_ is the effective size without background selection. In terms of the per base mutation rate *u* and recombination rate *ρ, U* = 2 *u l* and *R* = *ρ l*, where *l* is the length of the genomic region. We used values of *N*_e_ = 1333, *R* = 10^−8^×10^5^ = 10^−3^, *U* = 2×5×10^−8^×10^5^ = 10^−2^, *h* = 0.5, and varied the value of *s* (see Figure 1) in equation (2) to generate *N*_e,bg_ values. These *N*_e,bg_ values were then used as the effective population size *N*_e_ in equation (1) to generate the predicted fixation probabilities that are presented in Figures 1 and S2.

## Results

Background selection did not increase the range of cost/benefit ratios under which altruism is favored relative to Hamilton’s rule with *r* = 1/2; i.e., the fixation probability of the altruistic allele exceeded the neutral expectation when *c/b* < *r* (fixation probabilities are higher than the dashed line for *b* > 0.2 in Figure 1) and was less than the neutral expectation when *c/b* > *r* (fixation probabilities are lower than the dashed line for *b* < 0.2 in Figure 1). Instead, in accordance with the predictions from equations (1) and (2), background selection primarily changed the effective population size, which resulted in an increase in the altruistic allele’s fixation probability relative to no background selection when the allele was disfavored under Hamilton’s rule (compare intermediate *s* values to *s* = 0 and *s* = –1 for *b* < 0.2 in Figure 1) and in a decrease in its fixation probability when the allele was favored (compare intermediate *s* values to *s* = 0 and *s* = –1 for *b* > 0.2 in Figure 1). The effect was more extreme when there was stronger selection for or against the allele; for example, when the benefit of being helped was 0.19, fixation occurred rarely with background selection present and never without it (Figure 1). The allele was effectively neutral when the inclusive fitness effect (– *c* + *r b*) was 0, which occurred in two cases: when both the benefit and the cost were 0 and when the benefit and cost canceled one another, which occurred when the benefit *b* = 0.2, cost *c* = 0.1. In both of these cases, background selection doesn’t seem to affect fixation probabilities (Figure 1). These effects are consistent with background selection only affecting the fixation probability of the altruistic allele through a change in the effective population size *N*_e_. Further confirmation of this can be seen in Figure 1 where the predicted fixation probabilities assuming background selection only affects N_e_ are generally consistent with the simulation results.

A logistic regression found statistical support for the statements above. Using the simulation data where the cost of altruism is *c* = 0.1, we found that [1] the fixation probability increased strongly with the benefit of the altruistic allele, [2] the fixation probability increased modestly with the presence of background selection (i.e. *s* ≠ 0 and *s* ≠ –1), and [3] there was a significant negative interaction between these variables – that is, background selection increased the fixation probability when the benefit was small, and decreased the fixation probability when the benefit is large. Each result is highly significant (p < 10^−16^, for each predictor. See Supplementary Table S1 for model estimates and uncertainty in these estimates).

To further explore the effect of different recombination rates on the strength of background selection in our simulations, we ran simulations with both a higher and a lower recombination rate. As expected, varying recombination rate did not affect the cost/benefit ratios under which the altruistic allele was favored. Instead, the smaller recombination rate of 10^−12^ increased the effect of background selection and the larger recombination rate of 10^−4^ erased the effect of background selection (Supplementary Figure S2).

## Discussion

Research into the evolution of genes that affect altruism has largely ignored the genomic context of those genes, such as the possibility that linked selection might affect their evolutionary fates. Since background selection can increase genetic relatedness in the genomic regions where it occurs, we examined the effect of background selection on a genomic region that contains an altruistic allele. We used simulations where altruism was determined by a single allele and altruistic individuals could help their full siblings. We found that whether the altruistic allele was positively or negatively selected for was not affected by background selection, and could still be predicted by the classic condition for the evolution of altruism for family-structured populations, which is Hamilton’s rule. Instead, background selection seems to simply reduce the effective population size, *N*_e_. Stronger background selection, caused by weaker recombination or purifying selection in the genomic region, decreases *N*_e_ more. Decreased *N*_e_ weakens the effect of selection at the altruistic allele relative to genetic drift and shifts the fixation probability closer to the neutral value; thus, background selection in this scenario is increasing the fixation probability of deleterious alleles and decreasing the fixation probability of beneficial alleles, which is a well-known effect (Birky & Walsh, 1988; Charlesworth, 1994). Fixation probabilities calculated assuming background selection only affects *N*_e_ reasonably fit the simulation data and confirm that this assumption is relevant in this scenario.

After analyzing our results and finding that background selection did not affect whether the altruistic allele was under positive or negative selection, we went back to the foundational demographic assumptions in our model to see if we could explain this lack of an effect. Our model assumes a simple family-structured population where altruism occurs among full siblings. Inclusive fitness theory (Hamilton, 1964; Rousset, 2004) shows that the relatedness that matters for selection on a social trait is the relatedness between the individual expressing the social trait, the so-called “actor”, and the individual whose fitness is affected by the actor, the so-called “recipient”, at the time the trait is expressed (Taylor, 1990; Taylor & Frank, 1996). In the context of social interactions among full siblings, the baseline relatedness is naturally *r* = 1/2. The question is then how background selection affects genetic relatedness among siblings in this family-structured context. The original theory of background selection demonstrates how it affects relatedness among individuals randomly sampled from a population (Charlesworth et al., 1993; Hudson & Kaplan, 1995). However, background selection should have no effect on the relatedness between siblings within a family because selection against deleterious alleles acts only on adults and has no effect on the frequency of alleles within sibships. Thus, the sibling relatedness value of *r* = 1/2 is the correct value to use in Hamilton’s rule for this scenario. This reasoning suggests that background selection may have a more powerful effect on when altruism evolves in demographic scenarios where helping occurs between individuals whose relatedness can be affected by linked selection. One such scenario may be in spatially-structured populations when individuals help others in their local deme and background selection increases genetic relatedness above the level set by limited migration. However, Matthey-Doret and Whitlock (2019) show that the relationship between background selection and *F*_ST_ is complex since background selection can decrease both heterozygosity within demes and in the whole metapopulation and *F*_ST_ is a function of the ratio of these quantities. They found that background selection has little effect on the strength of relatedness measured by *F*_ST_ using parameter values drawn from empirical data on humans and sticklebacks. However, stronger background selection via higher deleterious mutation rates in larger regions with lower recombination rates and in populations with weaker migration rates than those tested by Matthey-Doret and Whitlock can significantly affect *F*_ST_ (Zeng & Corcoran, 2015; Matthey-Doret & Whitlock, 2019), which suggests that genomic variation in levels of background selection may be relevant to altruism in specific scenarios such as highly isolated populations with large inversions such as those found in supergenes (Schwander et al., 2014; Black & Shuker, 2019).

Support for the potential importance of the effect of linked selection on altruism comes from one of the few previous studies into how genomic context affects the evolution of social traits: Hammarlund et al. (2016) simulated mutations that confer a benefit in a new environment and are located on a non-recombining chromosome that also contains a social locus. Their results show that these beneficial mutations can favor the evolution of altruistic alleles in a spatially-structured population. While their scenario does involve linked selection, cooperation evolves in their scenario in a different way than the mechanism proposed here. Specifically, they assume that the size of each group in the population is positively related to the frequency of cooperation in that group, since more cooperative groups are likely to be more ecologically productive. Groups with more cooperators have more individuals, which increases the total chance that an adaptive mutation arises in those groups relative to groups with fewer cooperators. Given complete linkage between the beneficial loci and social locus, cooperative alleles can hitchhike alongside the beneficial alleles and rise in frequency with the beneficial alleles. This hitchhiking mechanism requires that the selection coefficient of a beneficial allele outweighs the cost of cooperation. In contrast, the hypothesis here is that linked selection due to background selection might have an effect because it increases genetic relatedness in regions linked to a social locus. Nevertheless, their result is intriguing and does suggest that selection on linked, “nonsocial”, mutations can impact the evolution of sociality given the right demographic circumstances.

Altruism is a complex behavior that can be subject to more complex evolutionary dynamics than non-social traits. However, our simulations show that altruism in a family-structured population is affected by background selection in the same way as simpler non-social traits are affected: background selection decreases the strength of selection for or against the linked trait. Nevertheless, altruism occurs in many more situations than portrayed by this single-population, sibling-helping-sibling model, and background selection may have a different effect in different scenarios. Further research into more complex population structures, different types of altruistic and cooperative interactions, and other kinds of linked selection may reveal more unique effects.

## Acknowledgements

LT was supported by an Undergraduate Research Opportunities Program (UROP) Fellowship from the University of Minnesota. JVC and DPE were supported by NSF CAREER DEB #1846260 and JVC by NSF DEB #1953223.

## Supplementary Figures

**Figure S1.**
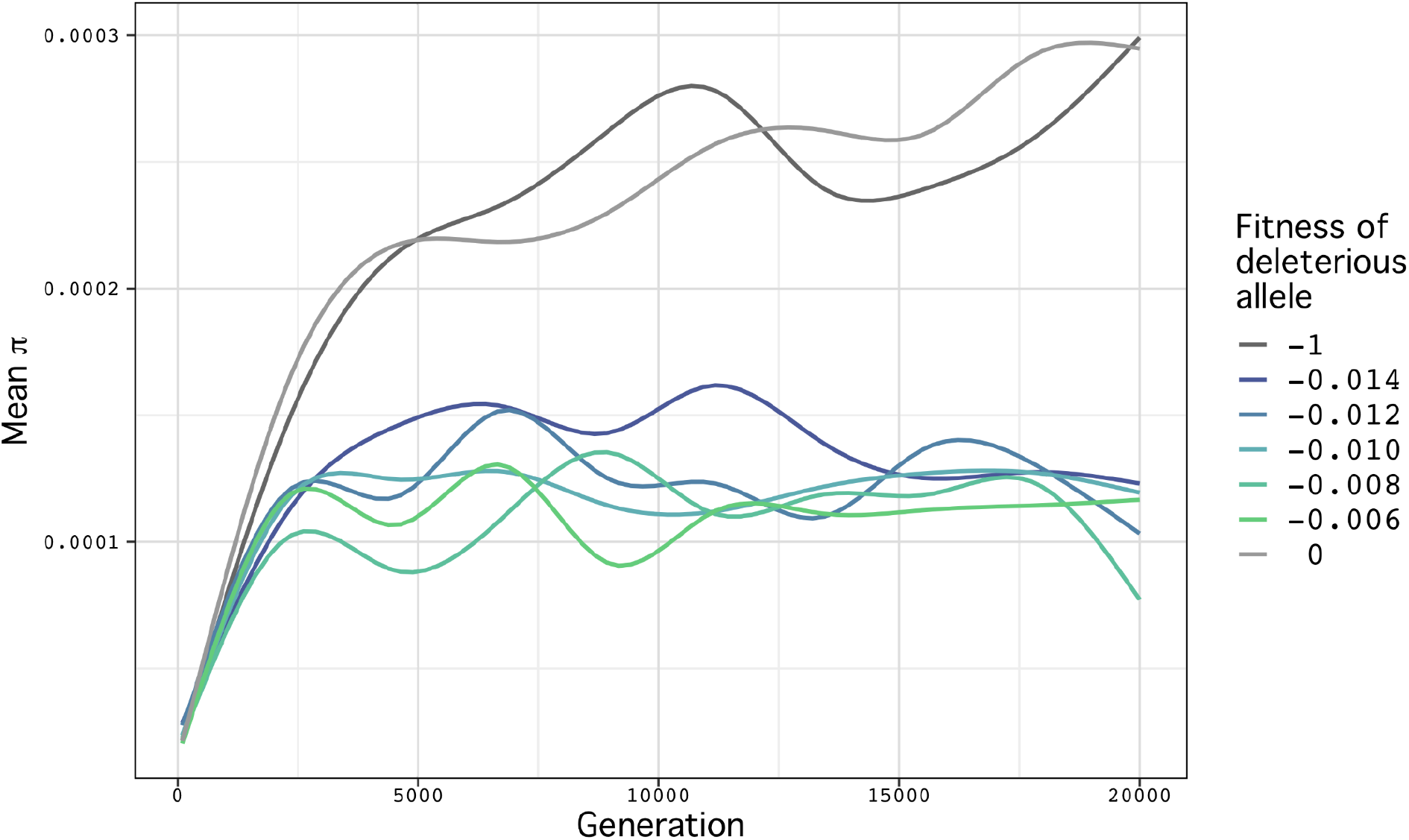
Genomic heterozygosity (*π*) over 20,000 generations in the presence and absence of background selection. 10 simulations were run for each fitness value for the detrimental mutations, and the mean *π* for each value is plotted. Background selection causes *π* to level off earlier and lower. For all fitness values, the greatest increase in *π* slows by generation 5000.

**Figure S2.**
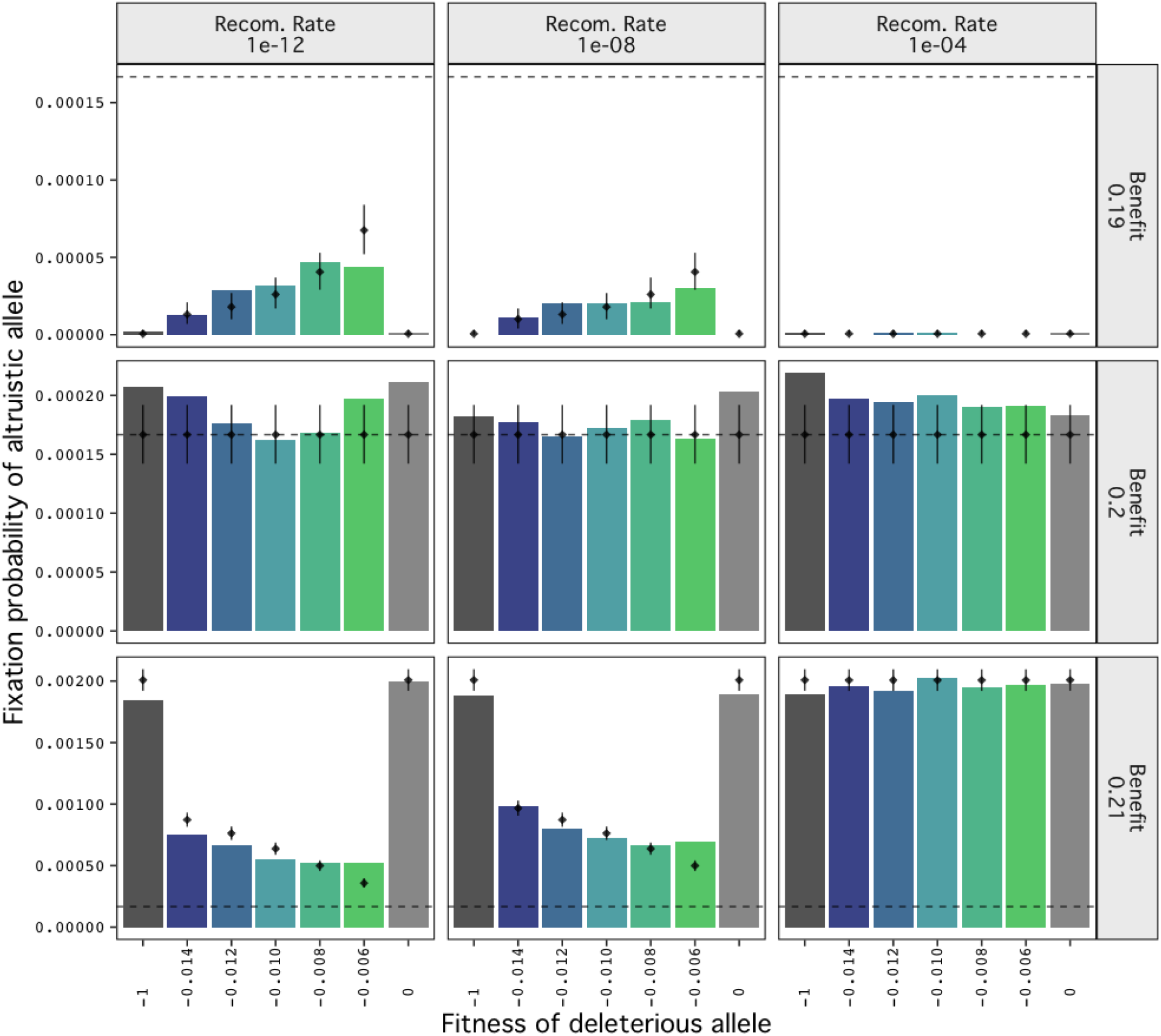
Varying recombination rate did not affect the cost/benefit ratios under which the altruistic allele is favored (*c*/*b* < 0.5 or *b* > 0.2 in the figure) and disfavored (*b* < 0.2 in the figure). Rather, the effect of background selection was stronger with a lower recombination rate of 10^−12^ while the higher recombination of 10^−4^ rate eliminated the effect of background selection. For the benefit *b* = 0.2, the allele was neutral at all recombination rates. The horizontal dashed line is the fixation probability of a neutral allele due to drift; vertical black lines indicate the 95% binomial confidence interval for the predicted fixation probabilities (see Methods for details).

**Table S1.** Results of our logistic regression model for the fixation probability of an altruistic allele. Cost is limited to 0.1. BGS is “no” if *s* = 0 or *s* = –1, and is “yes” otherwise.

| term | Estimate | Standard Error | z-value |
| --- | --- | --- | --- |
| (Intercept) | -59.07, | 0.572 | -103.21 |
| benefit | 251.523 | 2.764 | 91.01 |
| BGS | 18.932 | 0.686 | 27.584 |
| benefit x BGS | -94.384 | 3.329,-28.35 | -28.35 |

## References

Akçay, E. (2018). Collapse and rescue of cooperation in evolving dynamic networks. Nature Communications, 9(1), 2692. https://doi.org/10.1038/s41467-018-05130-7

Akçay, E., & Van Cleve, J. (2012). Behavioral responses in structured populations pave the way to group optimality. American Naturalist, 179(2), 257–269. https://doi.org/10.1086/663691

Allen, B., Lippner, G., Chen, Y.-T., Fotouhi, B., Momeni, N., Yau, S.-T., & Nowak, M. A. (2017). Evolutionary dynamics on any population structure. Nature, 544(7649), 227–230. https://doi.org/10.1038/nature21723

Axelrod, R., & Hamilton, W. D. (1981). The evolution of cooperation. Science, 211(4489), 1390–1396. https://doi.org/10.1126/science.7466396

Benton, T. G., & Foster, W. A. (1992). Altruistic housekeeping in a social aphid. Proceedings of the Royal Society of London. Series B: Biological Sciences, 247(1320), 199–202. https://doi.org/10.1098/rspb.1992.0029

Birky, C. W., & Walsh, J. B. (1988). Effects of linkage on rates of molecular evolution. Proceedings of the National Academyof Sciences, 85(17), 6414–6418. https://doi.org/10.1073/pnas.85.17.6414

Black, D., & Shuker, D. M. (2019). Supergenes. Current Biology, 29(13), R615–R617. https://doi.org/10.1016/j.cub.2019.05.024

Boomsma, J. J. (2009). Lifetime monogamy and the evolution of eusociality. Philosophical Transactions of the Royal Society B, 364(1533), 3191–3207. https://doi.org/10.1098/rstb.2009.0101

Bourke, A. F. G. (2014). Hamilton’s rule and the causes of social evolution. Philosophical Transactions of the Royal Society B, 369(1642), 20130362. https://doi.org/10.1098/rstb.2013.0362

Bourke, A. F. G., & Franks, N. R. (1995). Social evolution in ants (Vol. 62). Princeton University Press. https://doi.org/10.2307/j.ctvs32s3w

Burkart, J. M., Fehr, E., Efferson, C., & van Schaik, C. P. (2007). Other-regarding preferences in a non-human primate: Common marmosets provision food altruistically. Proceedings of the National Academyof Sciences of the United States of America, 104(50), 19762–19766. https://doi.org/10.1073/pnas.0710310104

Charlesworth, B. (1994). The effect of background selection against deleterious mutations on weakly selected, linked variants. Genetics Research, 63(03), 213–227. https://doi.org/10.1017/S0016672300032365

Charlesworth, B., Morgan, M. T., & Charlesworth, D. (1993). The effect of deleterious mutations on neutral molecular variation. Genetics, 134(4), 1289–1303. https://doi.org/10.1093/genetics/134.4.1289

Charlesworth, B., Nordborg, M., & Charlesworth, D. (1997). The effects of local selection, balanced polymorphism and background selection on equilibrium patterns of genetic diversity in subdivided populations. Genetics Research, 70(2), 155–174. http://journals.cambridge.org/article_S0016672397002954

Crow, J. F., & Kimura, M. (1970). An Introduction to Population Genetics Theory.

Harper & Row. Ewens, W. J. (2004). Mathematical Population Genetics. Springer. http://link.springer.com/book/10.1007/978-0-387-21822-9

Griesser, M., Drobniak, S. M., Nakagawa, S., & Botero, C. A. (2017). Family living sets the stage for cooperative breeding and ecological resilience in birds. PLoS Biology, 15(6), e2000483. https://doi.org/10.1371/journal.pbio.2000483

Haller, B. C., & Messer, P. W. (2019). SLiM 3: Forward genetic simulations beyond the Wright–Fisher model. Molecular Biology and Evolution, 36(3), 632–637. https://doi.org/10.1093/molbev/msy228

Hamilton, W. D. (1964). The genetical evolution of social behaviour. I. Journal of Theoretical Biology, 7(1), 1–16. https://doi.org/10.1016/0022-5193(64)90038-4

Hamilton, W. D. (1970). Selfish and spiteful behaviour in an evolutionary model. Nature, 228(5277), 1218–1220. https://doi.org/10.1038/2281218a0

Hamilton, W. D. (1972). Altruism and related phenomena, mainly in social insects. Annual Review of Ecology and Systematics, 3(1), 193–232. https://doi.org/10.1146/annurev.es.03.110172.001205

Hammarlund, S. P., Connelly, B. D., Dickinson, K. J., & Kerr, B. (2016). The evolution of cooperation by the Hankshaw effect. Evolution, 70(6), 1376–1385. https://doi.org/10.1111/evo.12928

Hu, X.-S., & He, F. (2005). Background selection and population differentiation. Journal of Theoretical Biology, 235(2), 207–219. https://doi.org/10.1016/j.jtbi.2005.01.004

Hudson, R. R., & Kaplan, N. L. (1995). Deleterious background selection with recombination. Genetics, 141(4), 1605–1617. https://doi.org/10.1093/genetics/141.4.1605

Kaplan, N. L., Hudson, R. R., & Langley, C. H. (1989). The “hitchhiking effect” revisited. Genetics, 123(4), 887–899. https://doi.org/10.1093/genetics/123.4.887

Krakauer, A. H. (2005). Kin selection and cooperative courtship in wild turkeys. Nature, 434(7029), 69–72. https://doi.org/10.1038/nature03325

Lehmann, L., & Keller, L. (2006). The evolution of cooperation and altruism–a general framework and a classification of models. Journal of Evolutionary Biology, 19(5), 1365–1376. https://doi.org/10.1111/j.1420-9101.2006.01119.x

Lehmann, L., & Rousset, F. (2014). The genetical theory of social behaviour. Philosophical Transactions of the Royal Society B, 369(1642), 20130357. https://doi.org/10.1098/rstb.2013.0357

Lukas, D., & Clutton-Brock, T. (2012). Cooperative breeding and monogamy in mammalian societies. Proceedings of the Royal Society B, 279(1736), 2151–2156. https://doi.org/10.1098/rspb.2011.2468

Matthey-Doret, R., & Whitlock, M. C. (2019). Background selection and FST: Consequences for detecting local adaptation. Molecular Ecology, 28(17), 3902–3914. https://doi.org/10.1111/mec.15197

Maynard Smith, J. (1974). The theory of games and the evolution of animal conflicts. Journal of Theoretical Biology, 47(1), 209–221. https://doi.org/10.1016/0022-5193(74)90110-6

Maynard Smith, J., & Haigh, J. (1974). The hitch-hiking effect of a favourable gene. Genetics Research, 23(01), 23–35. https://doi.org/10.1017/S0016672300014634

McAvoy, A., Allen, B., & Nowak, M. A. (2020). Social goods dilemmas in heterogeneous societies. Nature Human Behaviour, 4(8), 819–831. https://doi.org/10.1038/s41562-020-0881-2

Morgan, A. D., Quigley, B. J. Z., Brown, S. P., Buckling, A., & van Baalen, M. (2012). Selection on non-social traits limits the invasion of social cheats. Ecology Letters, 15(8), 841–846. https://doi.org/10.1111/j.1461-0248.2012.01805.x

Nadell, C. D., Drescher, K., & Foster, K. R. (2016). Spatial structure, cooperation and competition in biofilms. Nature Reviews Microbiology, 14(9), 589–600. https://doi.org/10.1038/nrmicro.2016.84

Noh, S., Geist, K. S., Tian, X., Strassmann, J. E., & Queller, D. C. (2018). Genetic signatures of microbial altruism and cheating in social amoebas in the wild. Proceedings of the National Academyof Sciences, 115(12), 3096–3101. https://doi.org/10.1073/pnas.1720324115

Ohtsuki, H., Hauert, C., Lieberman, E., & Nowak, M. A. (2006). A simple rule for the evolution of cooperation on graphs and social networks. Nature, 441(7092), 502–505. https://doi.org/10.1038/nature04605

Ostrowski, E. A. (2019). Enforcing Cooperation in the Social Amoebae. Current Biology, 29(11), R474–R484. https://doi.org/10.1016/j.cub.2019.04.022

Queller, D. C. (1985). Kinship, reciprocity and synergism in the evolution of social behavior. Nature, 318, 366–367. https://doi.org/10.1038/318366a0

Rankin, D. J. (2011). Kin selection and the evolution of sexual conflict. Journal of Evolutionary Biology, 24(1), 71–81. https://doi.org/10.1111/j.1420-9101.2010.02143.x

Rousset, F. (2004). Genetic Structure and Selection in Subdivided Populations (Vol. 40). Princeton University Press.

Rousset, F., & Billiard, S. (2000). A theoretical basis for measures of kin selection in subdivided populations: Finite populations and localized dispersal. Journal of Evolutionary Biology, 13(5), 814–825. https://doi.org/10.1046/j.1420-9101.2000.00219.x

Roze, D., & Rousset, F. (2004). The robustness of Hamilton’s rule with inbreeding and dominance: Kin selection and fixation probabilities under partial sib mating. American Naturalist, 164(2), 214–231. https://doi.org/10.1086/422202

Schwander, T., Libbrecht, R., & Keller, L. (2014). Supergenes and complex phenotypes. Current Biology, 24(7), R288–R294. https://doi.org/10.1016/j.cub.2014.01.056

Silk, J. B., & Boyd, R. (2010). From Grooming to Giving Blood: The Origins of Human Altruism. In P. M. Kappeler & J. Silk (Eds.), Mind the Gap: Tracing the Origins of Human Universals (pp. 223–244). Springer. https://doi.org/10.1007/978-3-642-02725-3_10

Slatkin, M., & Wiehe, T. (1998). Genetic hitch-hiking in a subdivided population. Genetics Research, 71(02), 155–160. https://doi.org/10.1017/S001667239800319X

Strassmann, J. E., Gilbert, O. M., & Queller, D. C. (2011). Kin discrimination and cooperation in microbes. Annual Review of Microbiology, 65(1), 349–367. https://doi.org/10.1146/annurev.micro.112408.134109

Strassmann, J. E., Zhu, Y., & Queller, D. C. (2000). Altruism and social cheating in the social amoeba Dictyostelium discoideum. Nature, 408(6815), 965–967. https://doi.org/10.1038/35050087

Tarnita, C. E., Ohtsuki, H., Antal, T., Fu, F., & Nowak, M. A. (2009). Strategy selection in structured populations. Journal of Theoretical Biology, 259(3), 570–581. https://doi.org/10.1016/j.jtbi.2009.03.035

Taylor, P. D. (1990). Allele-frequency change in a class-structured population. American Naturalist, 135(1), 95–106. https://doi.org/10.1086/285034

Taylor, P. D. (1992). Altruism in viscous populations –an inclusive fitness model. Evolutionary Ecology, 6(4), 352–356. https://doi.org/10.1007/BF02270971

Taylor, P. D., & Frank, S. A. (1996). How to make a kin selection model. Journal of Theoretical Biology, 180(1), 27–37. https://doi.org/10.1006/jtbi.1996.0075

Trivers, R. L. (1971). The evolution of reciprocal altruism. Quarterly Review of Biology, 46(1), 35–57. https://doi.org/10.1086/406755

Úbeda, F., & Gardner, A. (2010). A model for genomic imprinting in the social brain: Juveniles. Evolution, 64(9), 2587–2600. https://doi.org/10.1111/j.1558-5646.2010.01015.x

Uematsu, K., Kutsukake, M., Fukatsu, T., Shimada, M., & Shibao, H. (2010). Altruistic Colony Defense by Menopausal Female Insects. Current Biology, 20(13), 1182–1186. https://doi.org/10.1016/j.cub.2010.04.057

Van Cleve, J. (2015). Social evolution and genetic interactions in the short and long term. Theoretical Population Biology, 103, 2–26. https://doi.org/10.1016/j.tpb.2015.05.002

Van Cleve, J. (2017). Stags, hawks, and doves: Social evolution theory and individual variation in cooperation. Integrative and Comparative Biology, 57(3), 566–579. https://doi.org/10.1093/icb/icx071

Van Cleve, J. (2020). Building a synthetic basis for kin selection and evolutionary game theory using population genetics. Theoretical Population Biology, 133, 65–70. https://doi.org/10.1016/j.tpb.2020.03.001

Warneken, F., Hare, B., Melis, A., Hanus, D., & Tomasello, M. (2007). Spontaneous altruism by chimpanzees and young children. PLoS Biology, 5(7), e184. https://doi.org/10.1371/journal.pbio.0050184

Wolfram Research, Inc. (2021). Mathematica, Version 13.0.0. Wolfram Research, Inc. https://www.wolfram.com/mathematica

Wright, S. (1931). Evolution in Mendelian populations. Genetics, 16(2), 97–159. https://doi.org/10.1093/genetics/16.2.97

Wright, S. (1951). The genetical structure of populations. Annals of Eugenics, 15(1), 323–354. https://doi.org/10.1111/j.1469-1809.1949.tb02451.x

Zeng, K., & Corcoran, P. (2015). The effects of background and interference selection on patterns of genetic variation in subdivided populations. Genetics, 201(4), 1539–1554. https://doi.org/10.1534/genetics.115.178558

